# Stacking of anthocyanin related QTLs in the chromosome 10 and postulation of SmNac-like gene to be involve in anthocyanin biodhynthesis

**DOI:** 10.1101/2025.09.19.677371

**Authors:** Joaquin Gomis-Cebolla, Silvia Manrique, Andrea Arrones, Maria Dolores Toledo-Tolgar, Jose Luna, Virginia Baraja-Fonseca, Joan Sánchez-Pascual, Esther Gimeno-Páez, Mariola Plazas, Pietro Gramazio, Santiago Vilanova, Jaime Prohens

## Abstract

Anthocyanins in the fruit peel of photosensitive eggplants exhibit a different distribution pattern compared to the photo-insensitive ones. The latter exhibits a uniform anthocyanin content, whereas photosensitive eggplants lack anthocyanin accumulation in areas not exposed to light, such as under the calyx, or have lower concentrations in less-exposed areas. In the current research work, genetic analysis of F1 and F2 populations revealed that the photo-insensitive phenotype in eggplants follows an autosomal dominant inheritance with a 3:1 ratio, indicating that the photosensitive trait is regulated by a single dominant gene. To locate and narrow down the genomic region underlying photosensitivity, a segregating F2 population was used for bulked segregant analysis sequencing (BSA-seq) and compared with previous QTLs identified in previous developed eggplants populations (ILs and MAGIC population). The accumulation of QTLs at the end of chromosome 10 postulate that chromose region as a hot spot for anthocyanin related traits. In our population all the QTLs considered overlap between the genomic region 84,1-87,9 Mb. Moreover, no DNA mutations in the progenitors of the eggplant accessions used were found. A RNA-seq analysis of bagged photosensitive and photo- insensitive eggplants was performed, as a result we identified the *SmNAC1-like protein* gene as a promising gene to be involved in fruit photosensitivity trait. In the photo-insensitive accession (IVIA- 371) *SmNAC1-like protein* was depply repressed compare to the photosensitive accession (ASI-S-1). No consistent mutations in the coding sequences (CDS) of *SmNac-like protein* locus among all the different eggplants accessions used were found, suggesting that other layer of regulation maybe acting in our eggplant accessions. These findings provide new insight into the regulation of the molecular mechanisms of anthocyanin biosynthesis in eggplant as point out for the first time the possible role of NAC transcription factors in the anthocyanin biosynthesis in eggplant.

## INTRODUCTION

Eggplant (*Solanum melongena* L.) is a solanaceous crop with high economic value characterized by the presence of high concentrations of anthocyanins in the fruit peel of some cultivars. Anthocyanins are water-soluble pigments derived from anthocyanidins by the addition of sugars, responsible for red, blue or purple coloration of flowers, fruits and other plant organs (Takos et al., 2006 and Mattioli et al., 2020). Anthocyanins offer several benefits to plants, such as protection against UV radiation, extreme temperatures, and pathogen attacks (Lorenc-Kukula et al., 2005; Liu et al., 2013; Ren et al., 2014). In addition, anthocyanins contribute to human health, thanks to their strong antioxidant properties (Muñoz-Falcón et al., 2009; Speer et al., 2020).

In accordance with their role as photoprotectant pigments, light regulates the synthesis of anthocyanins in the fruit peel of eggplants. Previous research shows that light activates the anthocyanin biosynthetic pathway leading to the accumulation of anthocyanins in the fruit peel in a time-exposure manner (Jiang et al., 2016; Li et al., 2017; Sang et al., 2022; Li et al., 2023). However, there are some non-photosensitive eggplant varieties that are able to synthesize large amounts of anthocyanins independently of light conditions (Jiang et al., 2016 and Sang et al., 2022). An easy way to determine if an eggplant variety is able to synthetize anthocyanins independently from light is observing the presence of these pigments. under the calyx in the fruits. As these area is always shaded in photosensitive varieties it remains mostly anthocyanin-free (Li et al., 2017; Li et al., 2018 and Zhang et al., 2019). Consequently, breeding for photo-insensitive eggplant varieties is an effective method to obtain stable anthocyanin accumulation independently of environmental conditions. Therefore, several studies have focused on mapping the genes responsible for photoinsensitivity. These studies have identified different QTLs associated with non-photosensitivity phenotype across different chromosomes, likely due to the use of diverse eggplant materials and eggplant reference genomes (Qiao et al., 2022; He et al., 2022; Mangino et al., 2022; Luo et al., 2023a-2023b and You et al., 2023).

Several genes are involved in the biosynthesis of anthocyanin in eggplant (Jiang et al., 2016; Li et al., 2017; Li et al., 2018; Qiao et al., 2022; He et al., 2022; Mangino et al., 2022; Sang et al., 2022; Li et al., 2023; Luo et al., 2023ab and You et al., 2023). The so-called structural genes include enzyme- encoding genes such as *CHS*, *CHI*, *F3H*, *F3’H*, *F3*′*5*′*H*, *DFR* and *ANS* that catalyze the biosynthesis of anthocyanins (Jaakola et al., 2013 and Liu et al., 2018). While regulatory genes (RG) are the transcription factors (TFs) that control the expression of the structural genes (Li et al., 2017; Li et al., 2018; Qiao et al., 2022; He et al., 2022; Mangino et al., 2022; Zhou et al., 2022; Li et al., 2023; Luo et al., 2023a-2023b You et al., 2023). The transcription of structural genes is generally regulated by a complex formed by the interaction between a R2R3-MYB, a basic helix-loop-helix protein (bHLH) and WD repeat proteins (WD40), called MBW complex (<u>M</u>YB-<u>b</u>HLH-<u>W</u>D40) (MBW). Depending on the specific composition of the MBW complex, it can have a positive or negative effect over the transcription of structural genes (Ramsay et al., 2005; Yan et al., 2021). Since in normal conditions, anthocyanin biosynthesis is induced by light, the expression of RGs is, at its turn, controlled by different proposed keymaster genes (like HY5 or COP1,well-known regulators of light responses) widely spread among different species and participate in many metabolism pathways (Ramsay et al., 2005;Yan et al., 2021; Mankiota et al., 2024). Another well reported family of TFs known to regulate anthocyanin RGs that fall into our attention are NACs. The relationship between NACs and anthocyanins is that certain NAC transcription factors, such as *PpNAC25* (in peaches), *MdNAC52* (in apples) and *LcNAC13* (in lichi), act as positive regulators to promote anthocyanin biosynthesis (Jian et al ., 2019, Sun et al., 2019, Geng et al., 2022). Moreoover, overexpression of *PpNAC25* increase the production of anthocyanins in the fruit peel of peaches, the pigments responsible for the red colour in fruits such as peaches. These factors act by directing and activating specific genes involved in the anthocyanin biosynthesis pathway (Jian et al ., 2019, Sun et al., 2019, Geng et al., 2022)

In this study, we aimed to explore the genetic bases of anthocyanin photosensitivity in an F2 population derived from a cross between a photosensitive (ASI-S-1) and a non-photosensitive (IVIA- 371) *S. melongena* varieties. Through BSA-seq and RNA-seq, the TF *SmNAC-like protein* was postulated as a post promising gene to be involved anthocyanin photosensitivity. The *SmNac-like* was dow-regultaed in the non-photosensitive eggplant accession but no DNA sequence variants in the CDS consistent among the different eggplants accessions used was found. These results suggest that other layers of regulation may be interaction in the population used. The porpose of *SmNac1-like* as a loci to participate in eggplant photosensitive trait is useful to understand the genetic basis of the regulation of the anthocyanin pigmentation in the eggplant fruit peel, as well as for eggplant breeding.

## MATERIAL AND METHODS

### 1. Plant materials and cultivation conditions

The *S. melongena* accessions ASI-S-1 (anthocyanin photosensitive, non-striped, and with presence of chlorophyll pigmentation in the fruit peel) and IVIA-371 (anthocyanin non-photosensitive, striped, and no chlorophyll pigmentation in the fruit peel) were used as parents to develop an F2 population of 120 individuals segregating for the photosensitivity trait (PS), presence of fruit anthocyanin (FA), and chlorophyll content (CC) (Figure 1 Panels A and B). The PS trait was categorized as PUC (presence of purple pigmentation under the calyx, i.e., negative for PS) or puc (absence of purple pigmentation under the calyx, i.e., positive for PS). In addition to the eggplant accessions ASI-S-1 and IVIA-371, two sub-introgression lines (IL-69(6)5-12 and IL-69(6)5-8, were included in the study. These sub- lines were derived from a photo-insensitive *S. melongena* MEL5 accession that carries an introgression from the wild relative *S. incanum* on chromosome 10. Notably, these sub-lines exhibit a complete absence of anthocyanin content in the fruit peel (Figure S1). Additionally, the DH-ECAVI accession (photo-insensitive, with uniform anthocyanin pigmentation) was used, as well as H15 and AN-S-26 (photosensitive, exhibiting anthocyanin accumulation in the fruit peel when exposed to sunlight) (Mangino et al., 2022). Seeds of all the accessions were germinated in Petri dishes following the Ranil et al. (2015) protocol and transferred to seedling trays in a climatic chamber under a photoperiod 16 h light/ 8 h dark and temperature regime of 22 ± 2 °C. The plantlets were grown for four weeks (up to the fifth true leaf development stage) in a climatic chamber before acclimatization for 48 hours in a benched glasshouse. Plantlets were then transplanted to 15 L pots filled with coconut fiber and grown in a pollinator-free benched glasshouse of the Universitat Politècnica de València (UPV), Valencia, Spain. Plants were fertirrigated using a drip irrigation system and trained with vertical strings. Pruning was done manually to regulate vegetative growth and flowering. Phytosanitary treatments were performed when necessary.

**Figure 1.**
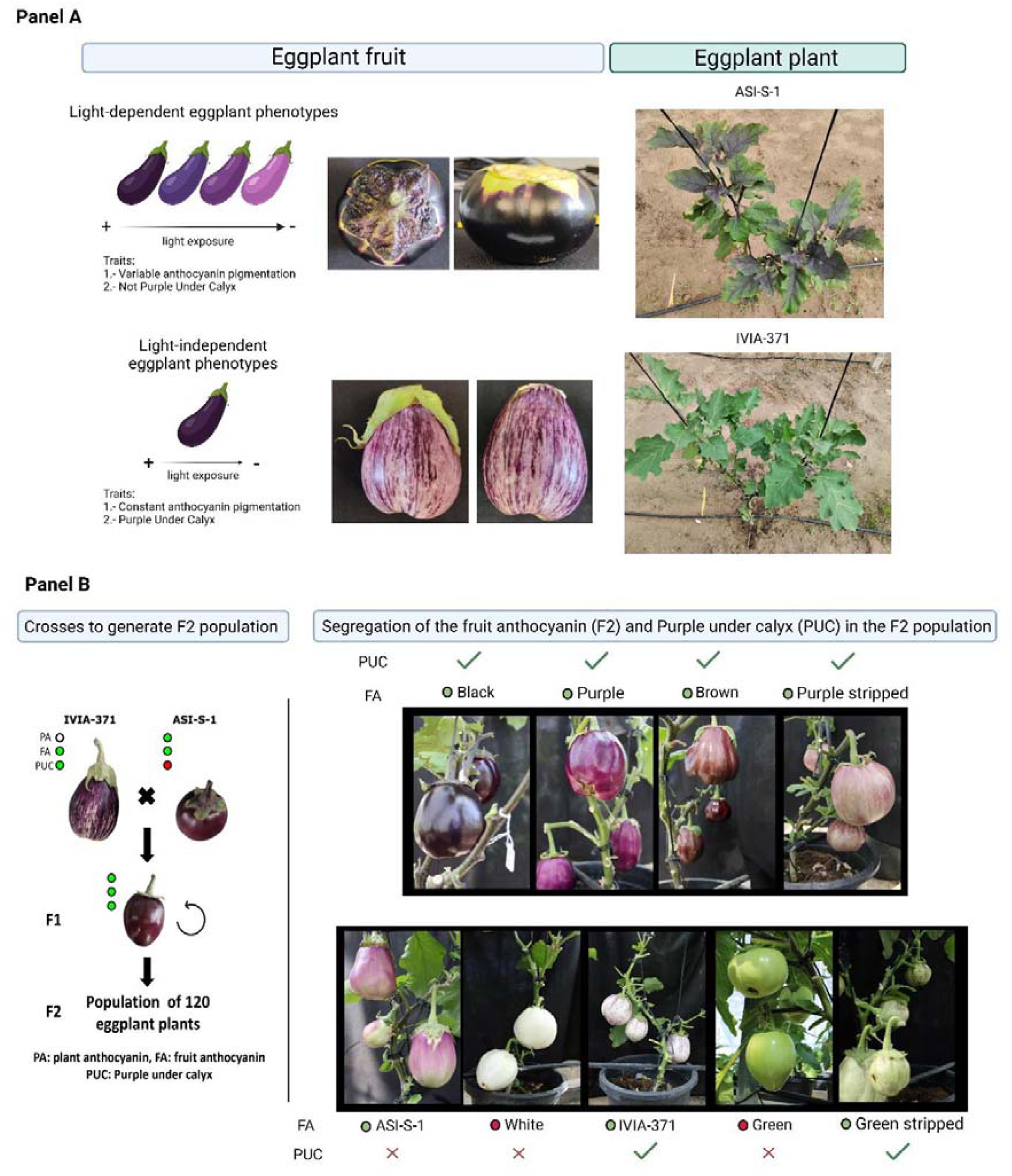
Phenotype of the *Solanum melongena* varieties photosensitive (ASI-S-1) and photoinsensitive (IVIA-371) and F2 population. Panel A: Phenotype of the eggplant fruit of ASI-S-1 and IVIA-371. Panel B: Crosses performed to develop the F2 population and the different F2 phenotypes obtained for fruit anthocyanin (FA), purple under calyx (PUC) and chlorophyll content (CC).

### 2. Generation of a F2 population and BSA-Seq analysis of PUC vs puc bulks

The F2 population obtained by the intraspecific cross between ASI-S-1 and IVIA-371 accessions was phenotyped for the PS and FA fruit traits. A Chi-square test was performed to assess the goodness-of-fit to a monogenic dominant inheritance pattern (3:1 segregation model). The F2 population was used for a BSA-Seq for the PS trait, which was scored in 94 plants out of the 120 of the total population, as 26 plants did not display FA. The PUC and puc bulks consisted of 15 F2 individuals. Young leaves were harvested from the entire F2 population (120 F2 individuals) as well as from six individual plants for each of the two parents. Their DNA was isolated, quantified, and tested for quality prior to select six plants from each of the parents and 30 individual F2 plants based on their different PS phenotype (PUC vs. puc). Genomic DNA was extracted using the silica matrix (SILEX) protocol (Vilanova et al., 2020) and checked for integrity by agarose electrophoresis (1% w/v). NanoDrop (Thermo Fisher Scientific, Waltham, MA, USA) ratios 260/280 and 260/230 were used to test the DNA quality, while its concentration was estimated with a Qubit 2.0 Fluorometer (Thermo Fisher Scientific, Waltham, MA, USA).

Pools for the PUC and puc phenotypes bulks were made by mixing in a equimolecular amount of DNA (quantified with Qubit 2.0 Fluorometer (Thermo Fisher Scientific) and AccuBlue ® Broad Range DNA quantitation kit (Biotium)) from each of selected F2 individual and shipped to Novogene (Novogene Europe, Cambridge, United Kingdom), where genomic libraries (PE 150, insert 350 bp) were constructed and sequenced. All raw reads were trimmed using the fastq-mcf tool from the Ea-utils package (Aronesty, 2013) with “q 30 -l 50” parameters, and the overall quality was checked using FastQC v.0.12.1 (Andrews et al., 2010). Clean reads were mapped against the 67/3 v3 (for QTL comparison and DNA variant analysis in reported eggplant populations positive for PS trait) and v.4.1 (for data cross-referencing with RNA-seq) eggplant reference genome (Barchi et al., 2019). To map the clean reads BWA-MEM v.0.7.17–r1188 was used with default parameters (Li and Durbin et al., 2009). The ΔSNP-index was estimated using the QTL-seq software v.2.2.3 (Takagi et al., 2013) from BAM files.

### 3. RNA-seq analysis of F2 population

RNA-seq analysis was conducted on bagged fruits from the F2 population to ascertain the gene expression profile of eggplant fruits with a positive and negative PS trait . Eggplant flowers from each plant were manually pollinated and kept in opaque paper bags for five days in the dark. The time lapse was selected, as it was found to be the shortest time for the appearance of anthocyanin pigmentation.

Subsequently, a set of eggplant fruit peels was gathered by cutting them with a surgical blade avoiding as much as possible to collect eggplant flesh, and subsequently frozen in liquid nitrogen before being kept in storage at -80°C. The eggplant fruit peels were grouped in three randomized replicates for each of PUC and puc phenotypes. For the PUC samples, each replicate consisted of four fruit peels. However, for the puc samples, due to the limitation in the number of fruits available, each replicate consisted of three samples. The RNA extractions were performed with RNAqueous kit and Ambion Plant RNA isolation aid reagent (Invitrogen™ AM1912 and AM6990) following the manufacturer instructions. The RNA integrity and quality were checked by 1.5% agarose gel and with the spectrometer Nanodrop 1000 (Thermo Fisher Scientific). RNA concentration was measured using a Qubit 2.0 Fluorometer (Thermo Fisher Scientific) and AccuBlue ® Broad Range RNA quantitation kit (Biotium). The samples with the highest values of 260/280 and 260/230 ratios were selected, normalized for their concentration, and randomly pooled.

RNA samples were shipped to Novogene Europe (Cambridge, UK) for library preparation and sequencing. Briefly, mRNA was purified from total RNA using poly-T oligo-attached magnetic beads. Then, mRNA was fragmented and the first strand cDNA was synthesized using random hexamer primers, followed by the second strand cDNA synthesis. After adapter ligation, fragments were size-selected, PCR-amplified, and purified. The library was checked with Qubit and real-time PCR for quantification and bioanalyzer for size distribution detection. Quantified libraries were pooled and sequenced on Illumina platforms (NovaSeq 6000), according to effective library concentration and data amount (9 GB of data). With regards to RNA-seq read processing, clean reads were obtained by removing reads containing adapter, reads containing ploy-N and low-quality reads from raw data through Novogene in-house perl scripts and Q20, Q30 and GC content were calculated. All the downstream analyses were based on clean, high-quality data (Table S1). Reference genome (*S. melongena* 67/3 v4.1) and gene model annotation files were downloaded from Sol Genomics Network (SGN) website. Index of the reference genome was built and paired-end clean reads were aligned to the reference genome using Hisat2 v2.0.5 (Zhang et al., 2021). This tool was selected for mapping as it can generate a database of splice junctions based on the gene model annotation file, giving a better mapping result than other non-splice mapping tools. Reads numbers mapped to each gene were counted using featureCounts v1.5.0-p3 (Liao et al., 2014). Then, FPKM was calculated based on the length of each gene and reads mapped to it.

Differential expression analysis was performed using the DESeq2 R package (1.20.0), which uses a statistical model based on the negative binomial distribution for determining differential expression (Love et al., 2014). The resulting P-values were adjusted using the Benjamini and Hochberg’s approach for controlling false discovery rate (Benjamini and Hochberg., 1995). Genes with an adjusted P-value ≤ 0.05 were assigned as differentially expressed. Finally, Gene Ontology (GO) enrichment analysis of differentially expressed genes (DEG) was implemented by the clusterProfiler R package (Yu et al., 2012; Wu et al., 2021; Xu et al., 2024), which corrects gene length bias. GO terms with corrected P-value < 0.05 were considered significantly enriched by differentially expressed genes. The clusterProfiler R package was used to test the statistical enrichment of differential expression genes in KEGG pathways database (http://www.genome.jp/kegg/).

### 4. Evaluation of DNA sequence variants in photo-sensitive and photo-insensitive cultivars from different eggplant population

To identify DNA sequence variants in the DEGs located inside the QTL identified at chromosome 10 in photo-in/sensitive eggplant cultivars, it were analyzed against v4.1 eggplant reference genome (Barchi et al., 2019). Firstly the QTL of chromosome 10 determined with the eggplant reference genome “67/3” v4.1 were compared against the QTL gotten with eggplant genome reference v3 with D-Genies (Cabanettes, Klopp., 2018). As a result, no major breaks and inversions were detected, indicating that the gene synteny remains constant at the genomic DNA region under consideration. Secondly of all the clean reads of ASI-S-1 and IVIA-371 obtained in a previously research (Mangino et al., 2022) were mapped against the eggplant reference genome “67/3” v3 with BWA-MEM and only the uniquely aligned reads were selected. Regarding the variant calling was carried out again, due that the reported sequence changes reported by genes Mangino et al. (2022) were not validated by Sanger Sequencing (Table S2). With regard the new variant calling filters for the PS trait in the F2 interspecific cross between ASI-S-1 and IVIA-371 were performed with FreeBayes (Garrison and Marth., 2012) with the following parameters: --min coverage 10, --limit coverage 60, and –min alternate fraction 0.2. The predicted effects of the sequence variants identified were evaluated with SnpEff (Cingolaniet et al., 2012). The DNA variants that affect the regulatory sequences (5’-UTR and 3’-UTR) and/or the CDS (i.e., frameshift, indels, splicing variants, etc) for the PS trait in homozygosis were classified as “high impact”. The Integrative Genomics Viewer (IGV) tool (Robinson et al., 2022) was used to validate the variants detected by the combination of FreeBayes + SnpEff softwares manually by visual inspection, with the following criteria: a sequence change was considered as a DNA variant if the mutation was present in all the reads with at leas a maq quality > 40 . To expand the number of eggplant cultivars evaluated, eggplants with PS trait (photo-insensitive: DH-ECAVI, photosensitive: ANS-S-26, and H15) previously reported (Mangino et al., 2022) were evaluated. Clean reads of the respective eggplant accessions were again mapped against the eggplant reference genome “67/3” v3 with BWA-MEM and only the uniquely aligned reads were selected. Then, the variant calling was performed with FreeBayes (Garrison and Marth., 2012) and the predicted effects of the DNA variants detected were evaluated with SnpEff (Cingolaniet et al., 2012). as previously.

## RESULTS

### 1. Identification and inheritance of a QTL linked to photoinsensitivity. 1.1.- Generation and inheritance of the scored traits in the F2 population

To determine the PS trait segregation pattern, an F2 population was produced by mating the *S. melongena* parents ASI-S-1 (puc, non-striped, and with presence of CC in the fruit peel) with IVIA- 371 (PUC, striped, and with absence of CC in the fruit peel). Regarding the presence of plant anthocyanin pigmentation in the parents, ASI-S-1 displayed anthocyanin in leaves and stem, meanwhile IVIA-371 only showed anthocyanin content in the stem but not in the leaves (Figure 1 Panel A). Regarding the F1, it displayed the PUC phenotype and uniform (non-striped), FA pigmentation, together with a background CC in the fruit peel. In the F2 generation, the PS, FA and CC traits segregated resulting in several phenotypic combinations (Figure 1 Panel B). for the three traits present in the eggplant cultivars used in this study. The segregation pattern of the PS trait showed an autosomal dominant inheritance (Table 1), being eggplants with PUC and non-striped phenotype the most abundant with a percentage of 78% (Figure 1). Plants producing fruits without FA (i.e., green or white fruits), were not scored for the PS phenotypes as the phenotype could not be determined (Figure 1 Panel B). These results suggested that the PS trait in our IVIA-371 x ASI-S-1 population was controlled each by one dominant gene. On the other hand, although the FA and CC traits segregate in the F2 population, they were not used in this study as they are traits genetically independent to the PS trait (data not shown).

**Table 1.**
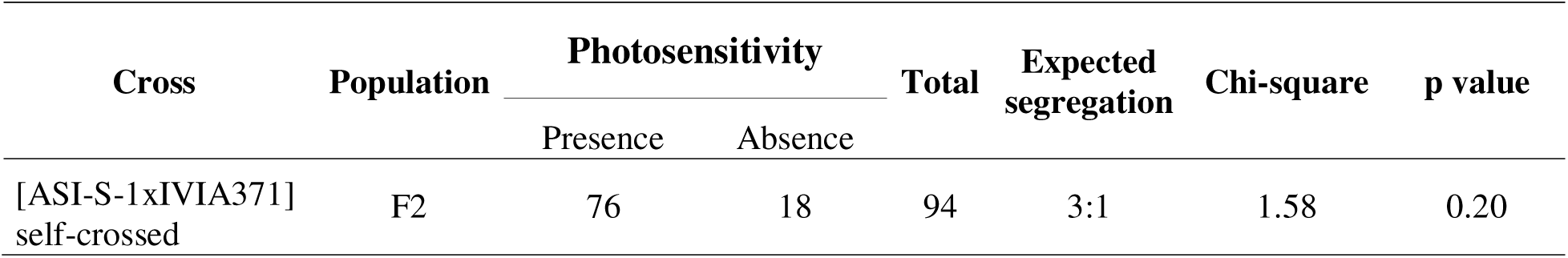
Genetic analysis of the segregation of photosensitivity in the F2 population from the cross of ASI-S-1 x IVIA-371. For photosensitivity, only the plants yielding fruits with anthocyanic pigmentation were used.

### 1.2. Identification of the genomic loci linked to the PS trait by bulk segregation analysis in the F2 population

To narrow down the genomic locus linked to the PS trait, two DNA pools from the IVIA-371 x ASI- S-1 F2 population (15 plants with the PUC phenotype and 15 with the puc phenotype) were sequenced. This sequencing yielded an average of 154 million 150-bp raw reads per bulk, with a mean coverage of 33.3x. Using ASI-S-1 as the reference sequence, the ΔSNP-index was calculated for 535,083 SNPs, getting a QTL on chromosome 10 between 62.9-69.7 Mb with a 99 % confidence interval (CI) for the PS trait. Similarly, when the ΔSNP-index was calculated for 562,169 SNPs using IVIA-371 as the reference sequence, a region between 63,4-67,5 Mb was identified on the same chromosome 10 with a 99 % CI (Figure 2, Figure S2, and Table S3). As a result, 102 genes were identified within the common candidate region (Table S4). Variant calling and prediction of the sequence variants effect were evaluated in both candidate regions of ASI-S-1 and IVIA-371. No DNA variants in the CDS (frameshift, indels, splicing variants, etc.) were detected.

**Figure 2.**
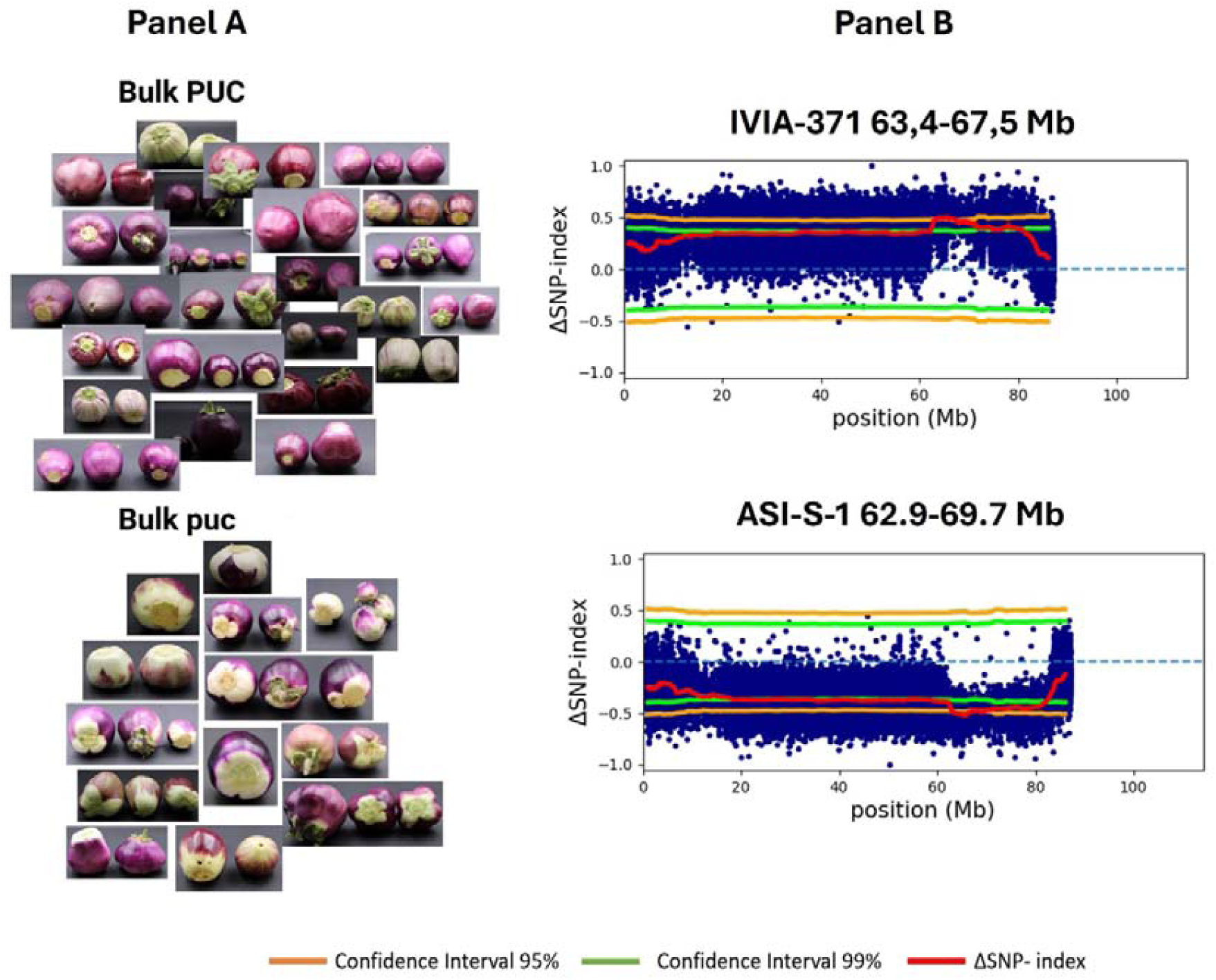
Results of the bulk segregation analysis in the F2 population of IVIA-371 (photoinsensitive) x ASI-S-1 (photosensitive). Panel A: Phenotype of the eggplants used to generate the bulks. Panel B: ΔSNP-index and the position of the respective QTL for IVIA- 371 and ASI-S-1, respectively.

### 2. Genetic evidences of QTL for photoinsensitivity in the F2 population and pevious developed eggplant populations

In order to determine the relevance of the QTL on chromosome 10 for the photosensitivity trait, it was compared with previous QTLs identified in different eggplant populations (Table 2). To be able to compare our data with the previous reported data (Muñoz-Falcón et al., 2009 and Mangino et al., 2022), the BSA-seq analysis was re-analyzed against the v3 of the eggplant genome reference (to use the same genome version in the comparison among the different populations). As a result of the re- analysis a QTL in the chromosome 10 for ASI-S-1 between 84,1-87,9 Mb with 99 % CI and a region between 83,5-97,7 Mb with 95 % CI for IVIA-371 were identified (Table S5, Table S6, Figure S2 and Figure S3).

**Table 2.**
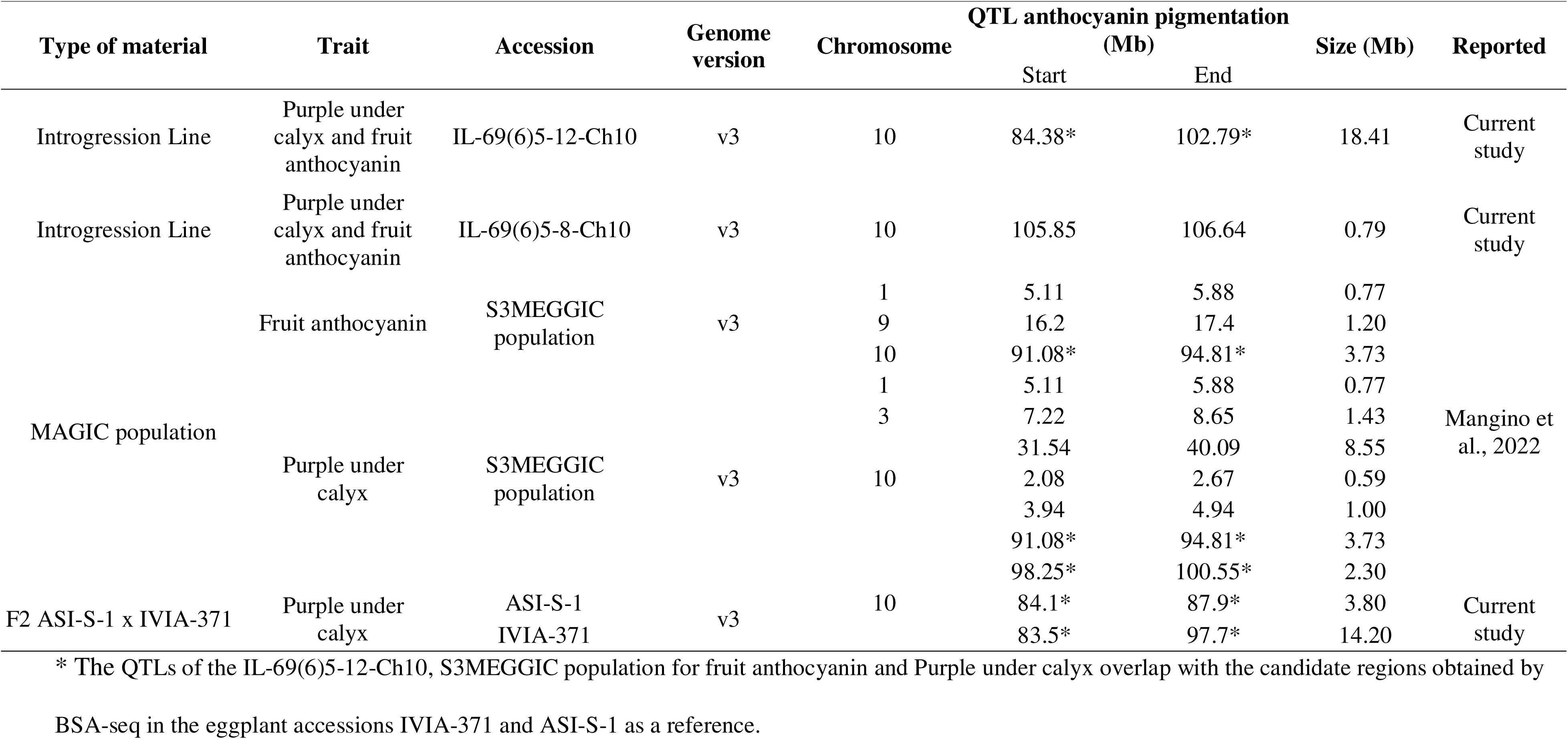
QTLs related with anthocyanin biosynthesis (fruit anthocyanin and purple under calyx traits) in different eggplant materials used in the current research and in Mangino et al. (2022), which used as parents of a MAGIC population some of the accessions used in this study.

Initially, two sub-introgression lines derived from introgressed DNA of *S. insanum* into *S. melongena* MEL5 were assessed. Specifically, the IL-69(6)5-12 and IL-69(6)5-8 contained an introgression on chromosome 10 at positions 84.4–102.8 Mb and 105.8–106.6 Mb, respectively, and both exhibited a complete loss of FA (Figure S1). Regarding the S3MEGGIC population, significant associated SNPs were identified at the end of chromosome 10 for both FA (91.1–94.8 Mb) and PUC (91.1–94.8 Mb and 98.2–100.5 Mb) (Table 2). Taking all the QTLs identified in the different populations together, suggest that the end of the chromosome 10 is a hotspot associated with anthocyanin-related traits. Six of the reported QTLs showed overlap (total or partial) along the different eggplant populations (Table 2). Of these six QTLs, two groups can be observed: the first one consists of four overlapping QTLs (IL-69(6)5-12-Ch10 84,4-102,8 Mb, FA 91.1-94.8 Mb, PUC 91.1-94.8 Mb, ASI-S-1 84,1-87,9 Mb, and IVIA-371 83,5-97,7 Mb); while the second one consists of two QTLs with partial overlap (IL- 69(6)5-12-Ch10 at 84,4-102,8 Mb and PUC 98.2-100.5 Mb). As a summary two common DNA regions across all the QTLs were identified (QTLs_ASI-S-1 84.1-87.9 (3.80 Mb) and PUC 98.2-100.5 (2.3 Mb)).

### 3 Differential expression analysis of bagged eggplants

The gene expression analysis between PUC and puc bagged eggplant fruits of the IVIA-371 x ASI-S- 1 F2 population was performed by RNA-seq of the fruit peel. The results indicated that from a total of 177 DEGs, 53 DEGs were up-regulated in the fruit peel of eggplants with PUC phenotype, while 124 DEGs were down-regulated in the fruit peel of eggplants with puc phenotype (Figure 3 and Table S7- 9). The 177 DEGs identified from the eggplant fruit peel for the PS trait could be grouped in four different subclusters (Table S9 and Figure S4). The subclusters 1 and 3 were made of the DEGs expressed mainly in IVIA-371 (PUC phenotype) with a total of 53 genes, that constituted the whole up-regulated DEGs identified by RNA-seq. The subclusters 2 and 4 contained 124 genes expressed totally in ASI-S-1 (puc phenotype), and constituted all the down-regulated DEGs reported for the eggplant fruit peel (Table S2 and Table S8). Concerning the functional and gene enrichment analysis of up- and down-regulated genes in eggplant fruit peel with PUC and puc phenotypes, few DEGS were identified due to the fruit peel characteristics. For the up-regulated DEGs expressed in IVIA- 371, the gene set showed an enrichment in KEGG terms related to flavonoid and phenylpropanoid biosynthesis (Table S7), mostly located in the subcluster 1 (Table S8) and in agreement with the FA presence (Figure 3 and Table S8). The following up-regulated genes in flavonoid biosynthesis were identified in IVIA-371: SMEL4.1_09g021180.1, SMEL4.1_05g002700.1, SMEL4.1_08g022350_1, SMEL4.1_02g020840_1, SMEL4.1_05g002760_1, SMEL4.1_05g018170_1, and SMEL4.1_12g016460_1. Regarding the GO annotation terms, an enrichment in terms like small molecule metabolic process, transferase activity of acyl and hexosyl groups, or coenzyme bonding and iron ion binding was observed. Those functions are consistent with the activity of the up-regulated genes identified by the KEGG terms.

**Figure 3.**
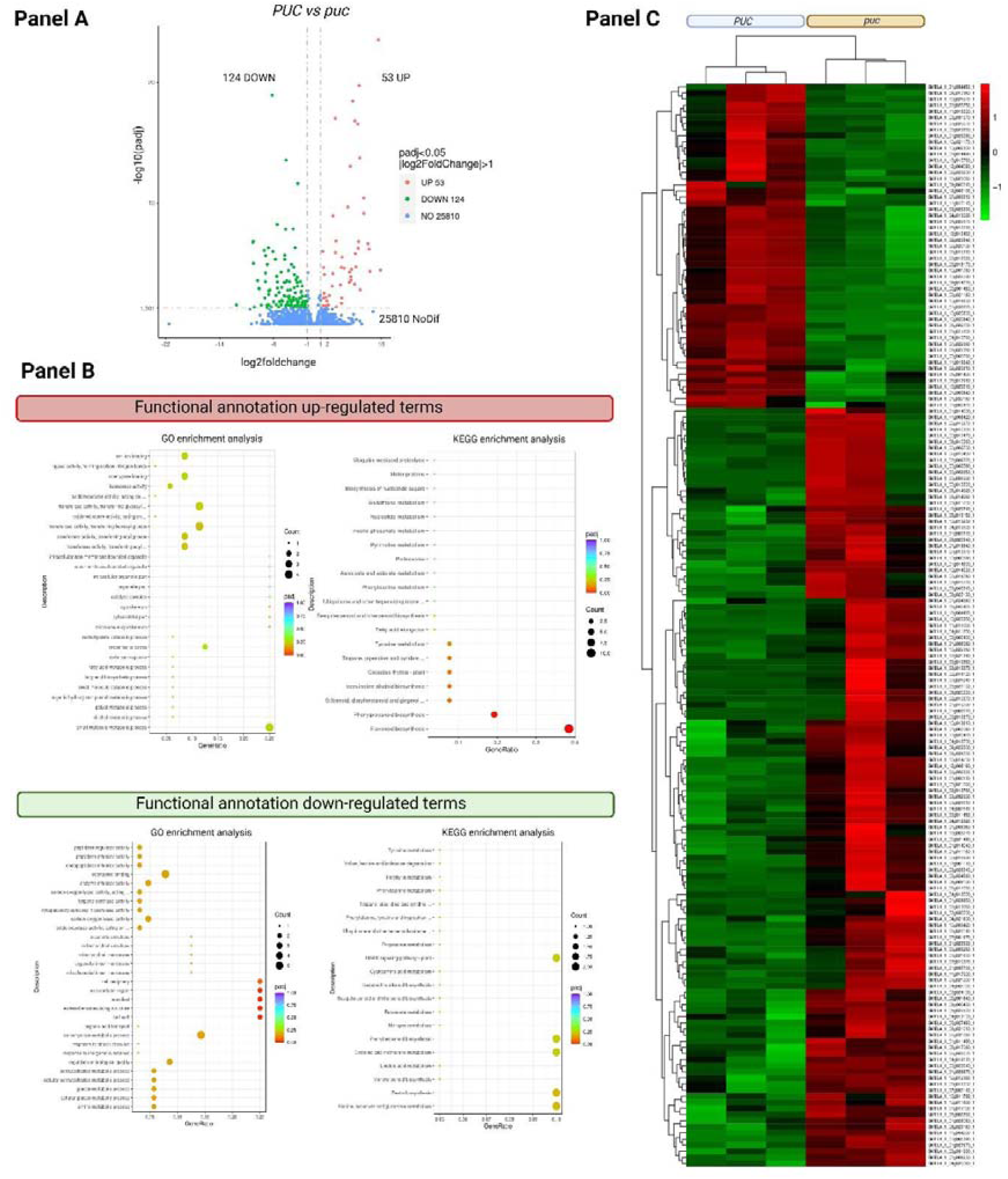
Differentially expressed genes (DEG) in early bagged photoinsensitive (IVIA-371) versus photosensitive (ASI-S-1) eggplants. Panel A: Vulcano plot of the up- and down- regulated of the fruit peel in early developed bagged eggplants. Panel B: Enrichment analysis of the up- down-regulated terms in GO and KEGG terms. Panel C: Clustering heatmap among the PUC vs puc and the DEG expressed in the eggplant fruit peel.

Concerning the down-regulated genes in IVIA-371 and, converserly, over-expressed in ASI-S-1, the gene set showed an enrichment in KEGG terms related to MAPK signaling pathway, phenylpropanoid biosynthesis, cysteine and methionine metabolism, zeatin and alanine biosynthesis, and aspartate and glutamate metabolism (Table S7). The enrichment in those KEGG terms in the MAPK signaling pathway and zeatin biosynthesis are involved in plant growth and fruit enlargement (Schäfer et al., 2015 and Jagodzik et al., 2018) in agreement with the fact that the eggplant fruits were growing. About the KEGG terms of cysteine and methionine metabolism plus alanine, aspartate and glutamate metabolism are reported to play a role in the generation of secondary metabolites involved in plant growth, fixing inorganic sulfur, precursors of several biosynthesis pathways, and plant protection against abiotic stress (Moormann et al., 2022). Finally, the accumulation of KEGG terms in phenylpropanoid biosynthesis indicated an accumulation of the biosynthesis precursors of flavonoids (Vogt, 2010; Dong et al., 2021 and Liu et al., 2021). With respect to the GO annotation terms an enrichment in terms like coenzyme binding, carbohydrate metabolic process, regulation of biological quality, or carbon-oxygen lyase activity was observed. Those functions are consistent with the activity of the up-regulated genes identified by the KEGG terms since they are involved in biosynthetic pathways.

### 4. Identification of genes up/down-regulated located in the QTL of chromosome 10 in photo-insensitive fruits in the dark conditions

To narrow down the number of putative candidate genes within the common candiate region of ASI- S-1 and IVIA-371 (102 genes), their expression levels (Table S9) were evaluated given that no DNA variant was identified see above – section 1.2). As a result, five genes were differentially expressed (*SMEL4.1_10g016630.1*, *SMEL4.1_10g014730.1*, *SMEL4.1_10g013610.1*, *SMEL4.1_10g014530.1*, and *SMEL4.1_10g019230.1*) (Table 3), while for the remaining 95 genes no changes in their expression were observed. The *SMEL4.1_10g016630.1*, which codifies for a *CHI-like protein*, an essential enzyme in the anthocyanin synthesis pathway, was up-regulated. This is in agreement with the fact that the PUC and puc pools showed anthocyanin content in the fruit peel (Figure 2). With regards the down-regulated genes (*SMEL4.1_10g014730.1*, *SMEL4.1_10g013610.1*, *SMEL4.1_10g014530.1*, and *SMEL4.1_10g019230.1), SMEL4.1_10g014730.1* and *SMEL4.1_10g014530.1* codify for a TF *NAC1-like protein* and a *RAD5B-like protein* (DNA repair protein), respectively. The remaining down-regulated genes encode for proteins of unknown function (Table 3). With the aim to known if the DNA mutations were present in the differentially expressed genes (Table 3) in DH-ECAVI (non-phosensitive), H15 and AN-S-26 (photosensitive). Firstly, all DNA sequences of the five genes differentially expressed identified in the v4.1 were located in the v3 of the eggplant refence genome. All of them were inside of the common DNA region of ASI-S-1 (84,1-87,9 Mb) and IVIA-371 (83,5-97,7 Mb) (Table S6). Secondly, the CDS were reevaluated in photosensitive (H15, ANS-S-26 and ASI-S-1) non-photosensitive (DH-ECAVI and IVIA-371) eggplant accessions. As a result, in the photoinsensitive and non-photosensitive eggplant no mutation in the CDS were consistent among all the eggplant accessions used

**Table 3.**
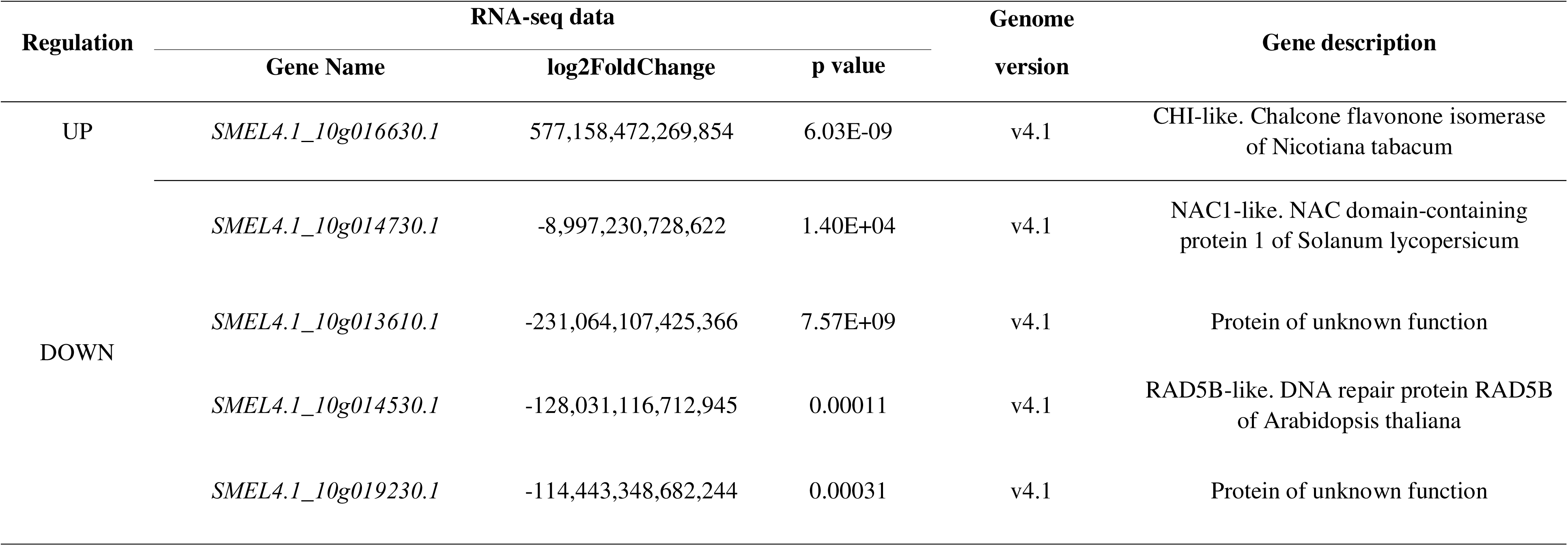
List of differentially expressed genes in the QTL of chromosome 10 delimited by bulk segregation analysis in the eggplant varieties IVIA-371 (photoinsensitive) and ASI-S-1 (photosensitive).

## DISCUSSION

Plant domestication in the Solanaceae family resulted in a great diversity of fruit shapes, sizes, and fruit colours. Despite this, modern breeding programs have generally favored eggplants with a uniform and intense dark and purple pigmentation in the fruit peel (Samuels et al., 2015; Wang et al., 2015 and Taher et al., 2017). In western markets, eggplants with black or purple peel are of higher economic value (Muñoz-Falcón et al., 2009). In our work, different eggplant populations were used to show that a genomic region of chromosome 10 is involved in antocyanin-related traits such as photosensitivy (PS) and fruit anthocyanin (FA). Anthocyanins have a strong antioxidant effect, so their high content in the fruit peel of eggplant improves nutrition. So, a research to determine the genetics basis of the PS is needed to set the basis of a new generation of eggplants cultivars with high content of secondary metabolites beneficial for human health. In the current research, the PS showed an inheritance in agreement with a monogenic dominant gene, following 3:1 ratio in an F2 generation, consistent with previous research data (Honda et al., 2014; He et al., 2022; Qiao et al., 2022; Luo et al., 2023ab). These eggplant accessions were selected because based on the corrected data reported by the GWAS study of the MAGIC population, neither of the two accessions presented mutations in the CDS of the candidate regions, suggesting that expression must have been altered.

Regarding the diversity of the fruit pigmentation of the F2 generation, this is probably due that more than one gene is contribuiting to the fruit peel pigmentation, in our F2 population at least three traits (PS, CC and FA) may interact to the final phenotype of the fruit peel pigmentation. The reported studies on the PS trait used commercial varieties of eggplant (Honda et al., 2012) or non-photosensitive eggplants varieties (He et al., 2022; Qiao et al., 2022; Luo et al., 2023ab) as parental donors for the PUC phenotype. Meanwhile for the puc phenotype present in photosensitive eggplants, it is recessive compared to the dominant allele of non-photosensitive eggplant varieties (Honda et al., 2012; He et al., 2022; Qiao et al., 2022; Luo et al., 2023a and 2023b). It should be noted that two commercial eggplant varieties with the PUC phenotype (IVIA-371) and (ASI-S-1) were used to produce the F2 population.

Contrarily to what occurs with other fruit colour traits such as the green fruit neeting (Arrones et al., 2022 and 2024; Toledo Garcia, et al., 2024), in which the phenotype present in the commercial eggplant varieties involves loss of gene function and recessive inheritance, the PS trait results in a dominant inheritance and a gain of function. That phenomenon is quite uncommon since domestication usually involves the loss of gene function or regulation (Lester and Hasan, 1991).

Concerning the localization of the reported QTLs (He et al., 2022; Qiao et al., 2022; Luo et al., 2023a and 2023b) for the PS trait at the end chromosome 10 using the eggplant genome 67/3 v3 as a reference across different eggplant accessions, is in agreement with our results. This supports that region as a promising target for fine-mapping or expression analysis to identify potential gene candidates. However, other QTLs for the PS trait have been reported in chromosome 1 and 3 (Mangino et al., 2022). Here, we confirmed the end chromosome 10, using v3 eggplant genome as a reference, as candidate region related with anthocyanin-related traits. To filter the number of candidates genes identified by the BSA-seq analysis using the eggplant genome version v4.1 (the QTL identified with v3 and v4.1 showed a 100% similarity), the list of genes was compared with the identified DEGs from the RNA-seq of an F2 population. From the 102 genes identified five of them were differentially expressed, with no mutations in their respective CDS in photo-insensitive (IVIA- 371 and DH-ECAVI) and photosensitive (ASI-S-1, H15 and AN-S-26) eggplant accessions. Specifically, out of the five genes differentially expressed, the homologous of the deeply repressed *SmNAC1-like* (SMEL4.1_10g014730.1) have been reported related to anthocyanin byosinthesis in *Prunus persica*, *Malus domestica* and *Litchi chinensis*. All these results together suggest *SmNAC1- like* gene as candidate gene controlling the PUC trait in our eggplant varieties.

As for the gene regulation of the anthocyanin biosynthetic pathway, previous studies suggest that light and cold induce anthocyanin biosynthesis from the moment of exposure until saturation through a highly dynamic range of TFs (Jiang et al., 2016; Li et al., 2017; Sang et al., 2022, Li et al., 2023 and Zhou et al., 2020). Specifically, *SmCry1*, *SmCry2*, *SmCry3* and *SmUVR8* are involved as photoreceptors in response to light. Meanwhile *SmCBF1*, *SmCBF2* and *SmCBF3* have been reported as sensors in response to low temperature. Both sets of genes activate the signaling pathway to produce anthocyanins (Li et al., 2017; Li et al., 2023 and Zhou et al., 2020). There is a consensus regarding the involvement of key structural genes in the downstream stages of anthocyanin biosynthesis pathway, including *SmCHI, SmF3H*, *Sm3GT*, *Sm5GT*, *SmDFR*, and *SmANS* (Jaakola et al., 2013 and Liu et al., 2018). For the intermediate steps, several TFs and their interactions have been proposed in different eggplant populations (R2R3-MYB transcription factor [*SmMYB113*, *SmMYB86*, *SmMYB75*, *SmMYB35*], basic helix–loop–helix [*SmbHLH1*, *SmbHLH13*], *SmHY5*, *SmTT8*, *SmGL3*, *SmTTG1*, *SmCIP7*, *SmGL2*, *SmGATA26*, *SmCOP1*, *SmSPA*, and *SmFTSH10*) (Jiang et al., 2016; Li et al., 2017; Li et al., 2018; Qiao et al., 2022; He et al., 2022; Mangino et al., 2022; Zhou et al., 2022; Li et al., 2023; Luo et al., 2023ab;and You et al., 2023). Specifically, any mutation in *SmMYB113* or pathway’s structural genes generate phenotypes without anthocyanin content (Mangino et al., 2022; Luo et al., 2023ab and You et al., 2023). Therefore, for the PS trait, the TFs proposed (*SmHY5*, *SmTT8*, *SmGL3*, *SmTTG1*, *SmCIP7*, *SmGL2*, *SmGATA26*) are in the first steps of the signaling pathway and always upstream of the *SmMYB113* (Mangino et al., 2022, Luo et al., 2023ab and You et al., 2023). This pattern also applied for *SmNAC1-like* where the homologous of *SmNAC1-like* in *Prunus persica* (*PpNAC1*), *Malus domestica* (*MdNAC1*) and *Lichi chinensis* (*LcNAC13)*) (Jiang et al., 2019, Liu et al., 2023 and Meng et al., 2023).

By using different eggplant materials, we have identified *SmNAC1-like* gene as a promising candidate for the PS trait, being highly repressed their expression in non-photosensitivity phenotypes. The eggplant PS trait has significant implications for fruit visual quality, as it results in uniform peel pigmentation, which is highly desirable for fruit commercialization. The study of this relevant trait could be of interest for future eggplant breeding projects with the aim of increasing the diversity of the fruit colour palette. In addition, due to a higher concentration of anthocyanin content and their effect as an antioxidant, *SmNAC1-like* could also be a valuable target for controlling the gene expression of the anthocyanin content in the fruit peel of the eggplants.

## ACKNOWLEDGEMENTS

This work has been funded by grants Juan de la Cierva (FJC2020-043194-I) funded by Agencia Estatal de Investigación and APOSTD-GVA (CIAPOS/2022/127) funded by Generalitat Valencia. Funding was also received from grant CIPROM/2021/020 funded by Conselleria d’Educació, Universitats i Ocupació of the Generalitat Valenciana, and from grants PID2021-128148OB-I00 and PID2024-160853OB-I00 funded by MCIN/AEI/10.13039/501100011033/ and by ERDF/EU. VB-F and J S-P are grateful to Generalitat Valenciana for grants INVEST/2022/146 and INVEST/2022/150, funded by European Union, Next Generation EU. JL is grateful to Spanish Ministerio de Ciencia, Innovación y Universidades for a predoctoral grant (FPU21/01590). SM would like to thank funding support from UPV and the Spanish Ministerio de Ciencia, Innovación y Universidades under the program María Zambrano funded by the European Union Next Generation-EU. PG is grateful to Spanish Ministerio de Ciencia, Innovación y Universidades for a post-doctoral grant (RYC2021–031.999-I) funded by MCIN/AEI /10.13039/501.100.011.033 and the European Union through NextGenerationEU/PRTR.

## CONTRIBUTIONS

JG-C, SM, PG, SV, MP and JP conceived the idea. JG-C and SM supervised and designed the manuscript. JG-C, SM, AA and MDT-T developed the F2 population for BSA-seq and RNA-seq. EG-P developed the introgression lines. JG-C, SM, JL, JS-P, MDT-T performed DNA and RNA extraction. JG-C wrote the manuscript. All the authors reviewed and edited the manuscript.

## DATA AVAILABILITY STATEMENT

The data presented in the study are deposited in the NCBI SRA repository, BioProject IDs for BSA-seq are the following: F2 photo-insensitive, SRS26640024, F2 photosensitive SRS26640023, ASI -S-1 SRS26640022 and IVIA-371 SRS26640021. With regards theRNA-se analysis are under the BioProject IDs: PUCR replicate 1 SRR35685429, PUCR replicate 2 SRR35685426, PUC replicate 3 SRR35685424, puc replicate 1 SRR35685427, puc replicate 2 SRR35685425, puc replicate 3 SRR35685428.

## GENERATIVE AI STATEMENT

The authors declare that no Gen AI was used in the creation if this manuscript or in any of the figure and table attached to the manuscript.

## SUPPLEMENTARY MATERIALS

**Table S1.** RNA quality data of the samples used in the RNA-seq data of PUC and puc.

**Table S2.** Evaluation of the gene related with anthocyanin biosynthesis in the QTLs determined by Mangino et al., 2022.

**Table S3**. Results of bulk segregation analysis in ASI-S-1 and IVIA-371 with a confidence interval of 95% and 99%, respectively. Eggplant genome reference v4.1

**Table S4.** List of genes in the candidate region acotated by BSA-seq in eggplant varieties ASI-S-1 and IVIA-371. Eggplant genome reference v4.1

**Table S5.** Results of bulk segregation analysis in ASI-S-1 and IVIA-371 with a confidence interval of 95% and 99%, respectively. Eggplant genome reference v3

**Table S6.** List of genes in the candidate region acotated by BSA-seq in eggplant varieties ASI-S-1 and IVIA-371. Eggplant genome reference v3

**Table S7.** List of differentially expressed genes in the eggplant varieties IVIA-371 (PUC positive trait) versus ASI-S-1 (puc negative trait).

**Table S8.** List of the subcluster identified in the heatmap of the differentially expressed genes of light-independent versus light-dependent eggplant varieties.

**Table S9**. List of the enrichment analysis of the GO and KEGG terms of the up- and down- regulated genes in PUC versus puc eggplant fruit peel

---

**Figure S1.**
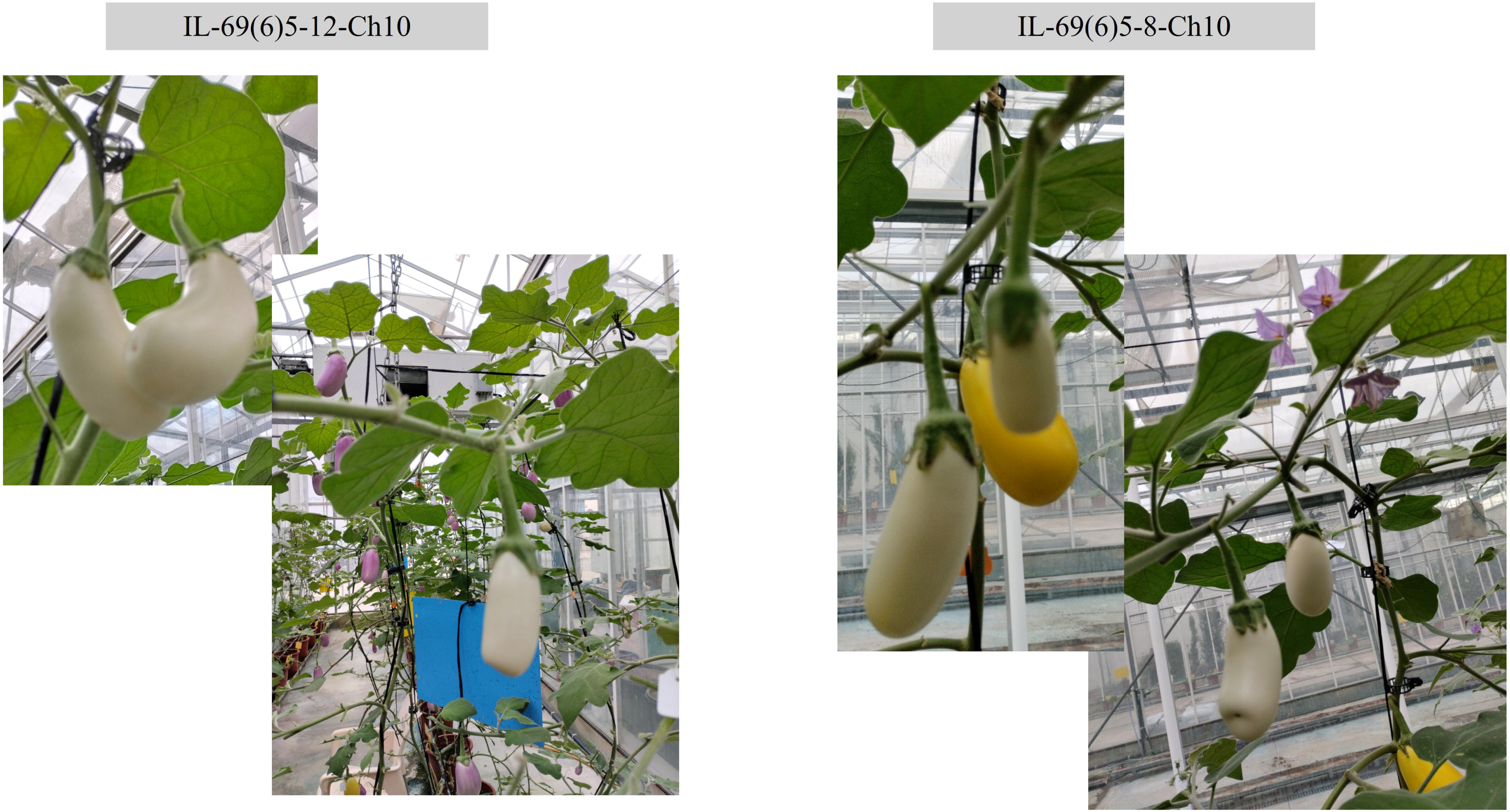
Phenotype of the two sub-introgression lines (IL-69(6)5-12-Ch10 and IL-69(6)5-8- Ch10) derived from the introgression line IL-M5-19.

**Figure S2.**
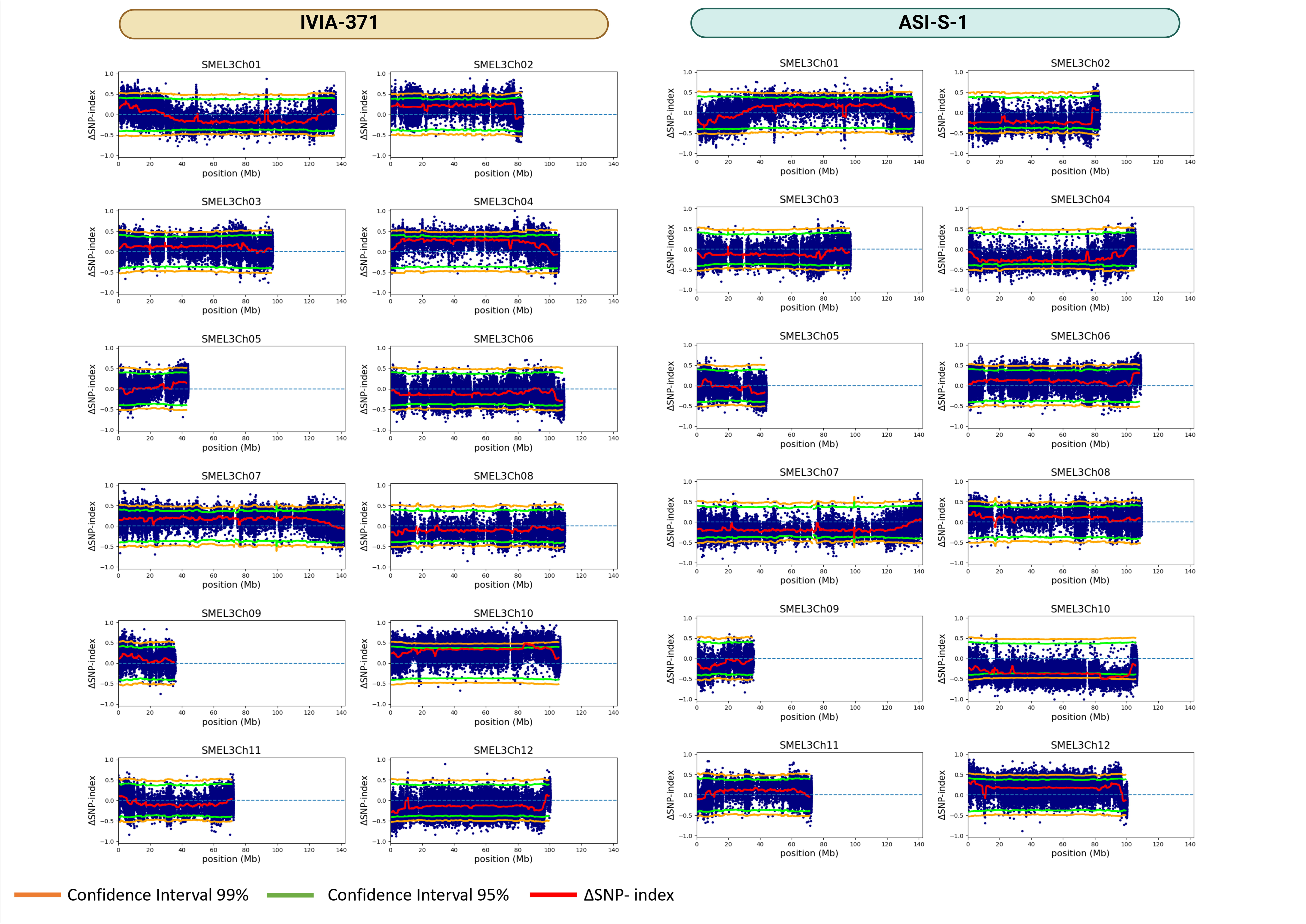
Delta of SNP-Index of the bulk segregation analysis of light-dependent and light- independent eggplant varieties

**Figure S3.**
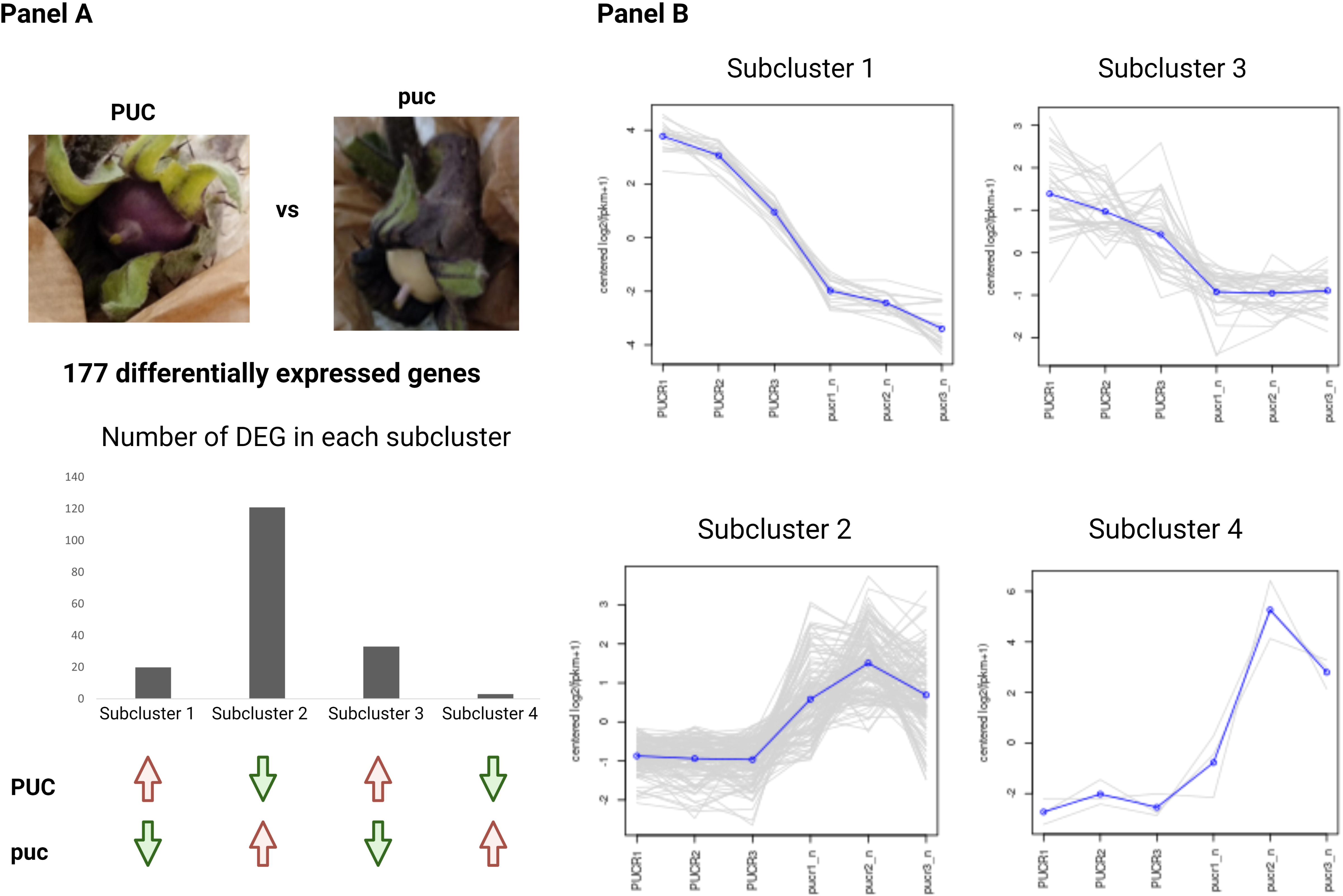
Number of differentially expressed genes (DEG) in the four subclusters identified in the clustering heatmap PUC vs puc. The DEG in the subcluster 1 and 3 were up-regulated in the light-independent (IVIA-371) eggplant (Panel A and B) meanwhile the DEG in the subcluster 2 and 4 were up-regulated in the light-dependent (ASI-S-1) eggplant.

